# Voltage dynamics of dendritic integration and back-propagation *in vivo*

**DOI:** 10.1101/2023.05.25.542363

**Authors:** J. David Wong-Campos, Pojeong Park, Hunter Davis, Yitong Qi, He Tian, Daniel G. Itkis, Doyeon Kim, Jonathan B. Grimm, Sarah E. Plutkis, Luke Lavis, Adam E. Cohen

## Abstract

Neurons integrate synaptic inputs within their dendrites and produce spiking outputs, which then propagate down the axon and back into the dendrites where they contribute to plasticity. Mapping the voltage dynamics in dendritic arbors of live animals is crucial for understanding neuronal computation and plasticity rules. Here we combine patterned channelrhodopsin activation with dual-plane structured illumination voltage imaging, for simultaneous perturbation and monitoring of dendritic and somatic voltage in Layer 2/3 pyramidal neurons in anesthetized and awake mice. We examined the integration of synaptic inputs and compared the dynamics of optogenetically evoked, spontaneous, and sensory-evoked back-propagating action potentials (bAPs). Our measurements revealed a broadly shared membrane voltage throughout the dendritic arbor, and few signatures of electrical compartmentalization among synaptic inputs. However, we observed spike rate acceleration-dependent propagation of bAPs into distal dendrites. We propose that this dendritic filtering of bAPs may play a critical role in activity-dependent plasticity.

## Introduction

Neuronal information processing is governed by a complex interplay among diverse voltage-dependent ion channels distributed throughout the dendritic tree, soma, and axon ^1,2,3,4^. Dendrites perform two qualitatively distinct information-processing tasks: they integrate synaptic inputs to determine spike times; and they integrate spikes and synaptic inputs to mediate synaptic plasticity ^5^. Broadly, these processes involve information flow in opposite directions: from synapse to soma, and from soma to synapse; yet both processes interact with the same set of highly nonlinear dendritic ion channels, and therefore must follow the same local excitation rules. A fundamental mystery in neurobiology is how the same set of dendritic channels mediates these two distinct tasks.

Experiments in acute brain slices have characterized many of the dendritic ion channels and their associated bioelectrical excitations ^6^. While slice preparations provide valuable insights, they cannot reveal the dynamics of information processing in a living organism, where the electrotonic status of the dendrites and their ion channels may be substantially different from slices, and where the spatial and temporal input patterns are, in general, unknown and dependent on brain state. Previous *in vivo* investigations of dendritic physiology ^7^ have primarily employed calcium imaging ^8,9,10,11^, and patch-clamp ^12,13,14,15,16^ techniques on single compartments, but it has not yet been possible to record voltage in multiple dendrites and soma *in vivo* at the same time, a necessary step toward mapping sub-cellular information flow.

To explore the voltage dynamics of dendritic arbors, we delivered a Cre-dependent Optopatch construct via *in utero* electroporation to drive expression in Layer 2/3 cortical neurons in the barrel cortex of adult mice. We combined a chemigenetic voltage indicator, Voltron2 ^17^, with a blue shifted channelrhodopsin (CheRiff) ^18^ linked to a fluorescent protein (eYFP) for visualization and a Lucy-Rho ^19^ sequence for improved membrane trafficking (**Fig. 1a**). The home-built imaging system contained two digital micromirror devices (DMDs) for independent dynamic targeting of blue light (for optogenetic stimulation) and orange light (for structured illumination voltage imaging). In the imaging path, we acquired simultaneous recordings from the soma and apical dendrites by splitting the fluorescence between two cameras whose focal planes could be adjusted independently (**Fig. 1b**). We designed our optical system and imaging headplate to minimize optical aberrations (Methods, **Fig. S1–S3**). Custom instrument-control software ensured microsecond-precision synchrony and sub-pixel spatial registration between both cameras and both DMDs. Post-processing analysis corrected for in-plane and axial motion, blood flow, and photobleaching (Methods, **Figs. S4–S7**). Images were acquired at either 500 or 1,000 Hz (Methods).

**Fig. 1.**
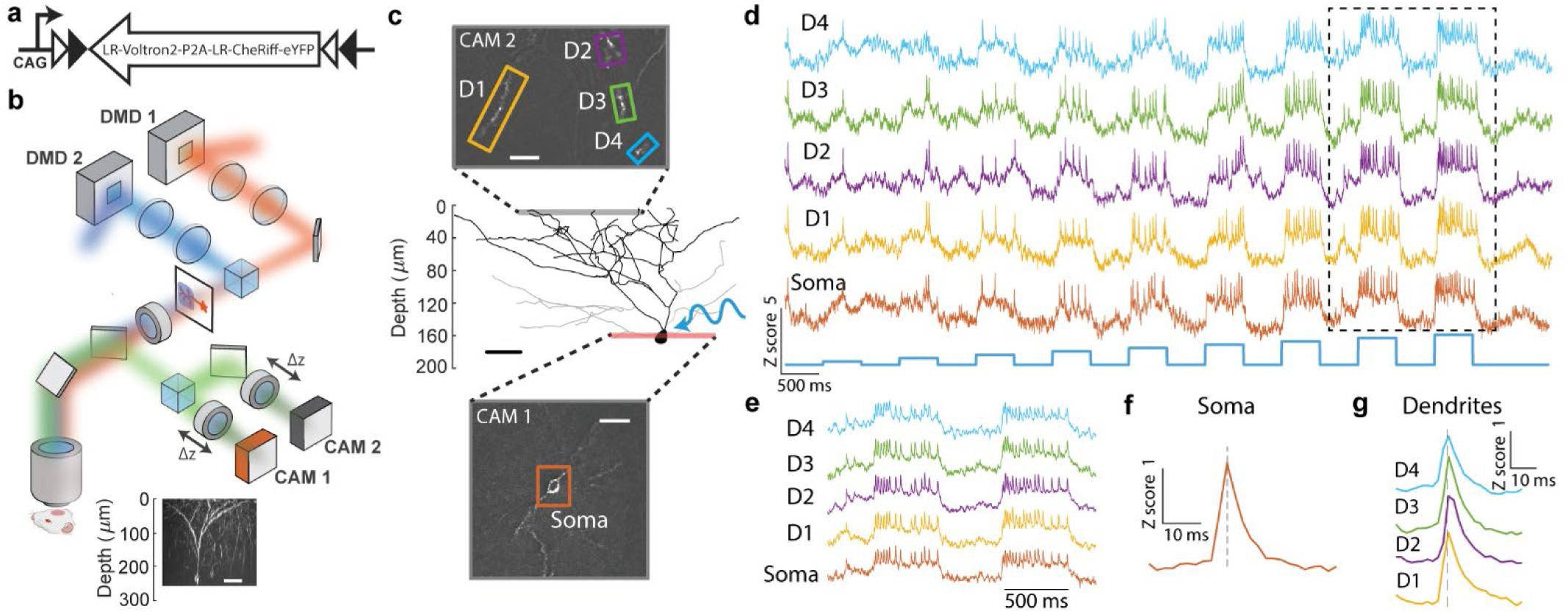
Dendritic voltage mapping and optogenetic stimulation *in vivo*. **a**, Construct for expressing Optopatch in cortical neurons, comprising a chemigenetic voltage indicator, Voltron2, and a blue-shifted channelrhodopsin, CheRiff. White and black triangles depict the lox2272 and loxP recombination sites, respectively. **b**, Instrument for targeted channelrhodopsin stimulation and structured illumination voltage imaging in two focal planes simultaneously. Inset: side projection of a single L2/3 neuron taken with a 1-photon HiLo z-stack (Methods). Scale bar 50 μm. **c**, Reconstructed morphology of a single neuron, with simultaneous recordings at the soma and at four apical dendrites. Scale bar 30 μm. **d**, Optogenetic stimulation with pulses of increasing blue light intensity evoked action potentials with increasing spike rates and corresponding bAPs in the dendrites. **e**, Close-up of boxed region in (d) showing correspondence between somatic and dendritic events. **f-g** Spike triggered average (STA) of fluorescence at the soma (f) and dendrites (g), triggered off the spike times at the soma. The dendritic events peaked after the somatic spike (dashed line), as expected for back-propagation.

By combining a red-shifted voltage indicator, very sparse expression, and structured illumination microscopy (HiLo) ^20^ (Methods) we could map and record neuronal compartments to a depth of 200-300 μm. **Figure 1c** shows a representative HiLo side projection of a single neuron with one somatic and four apical dendrite recording sites. We stimulated the neuron with 0.5 s pulses of widefield blue light at increasing intensity and simultaneously recorded from the soma and four dendrites (**Fig. 1c,d**). We observed robust blue light-evoked spiking in the soma, and bAPs in the dendrites. We calculated average bAP waveforms in the dendrites, triggered off the spike peaks at the soma, and confirmed that the soma spiked first, as expected for bAPs (**Fig. 1e,g**).

### Mapping bAP propagation

We mapped the propagation of bAPs in anesthetized mice by keeping CAM 1 focused at the soma and moving the field of view and focus of CAM 2 across different branches of the dendritic arbor. For each CAM 2 field of view, we evoked a spike train at the soma (40 pulses, 8 ms duration, 5 Hz) and recorded the spikes at the soma and the bAPs in the dendrites (**Fig. 2a,b**). For these optogenetically evoked single spikes, we did not observe trial-to-trial variation in the bAP amplitude beyond baseline noise fluctuations at any of the recording sites (see e.g., **Fig 2c**), indicating that bAP failures were infrequent under these conditions. We used a background-robust estimate of ΔF/F to ensure that signals were proportional to local voltage (Methods), and then calculated the mean bAP waveforms at each recording site (**Fig. 2d**), triggered by the peak timing at the soma. From the bAP waveforms we extracted the bAP amplitude and conduction delay.

**Fig. 2.**
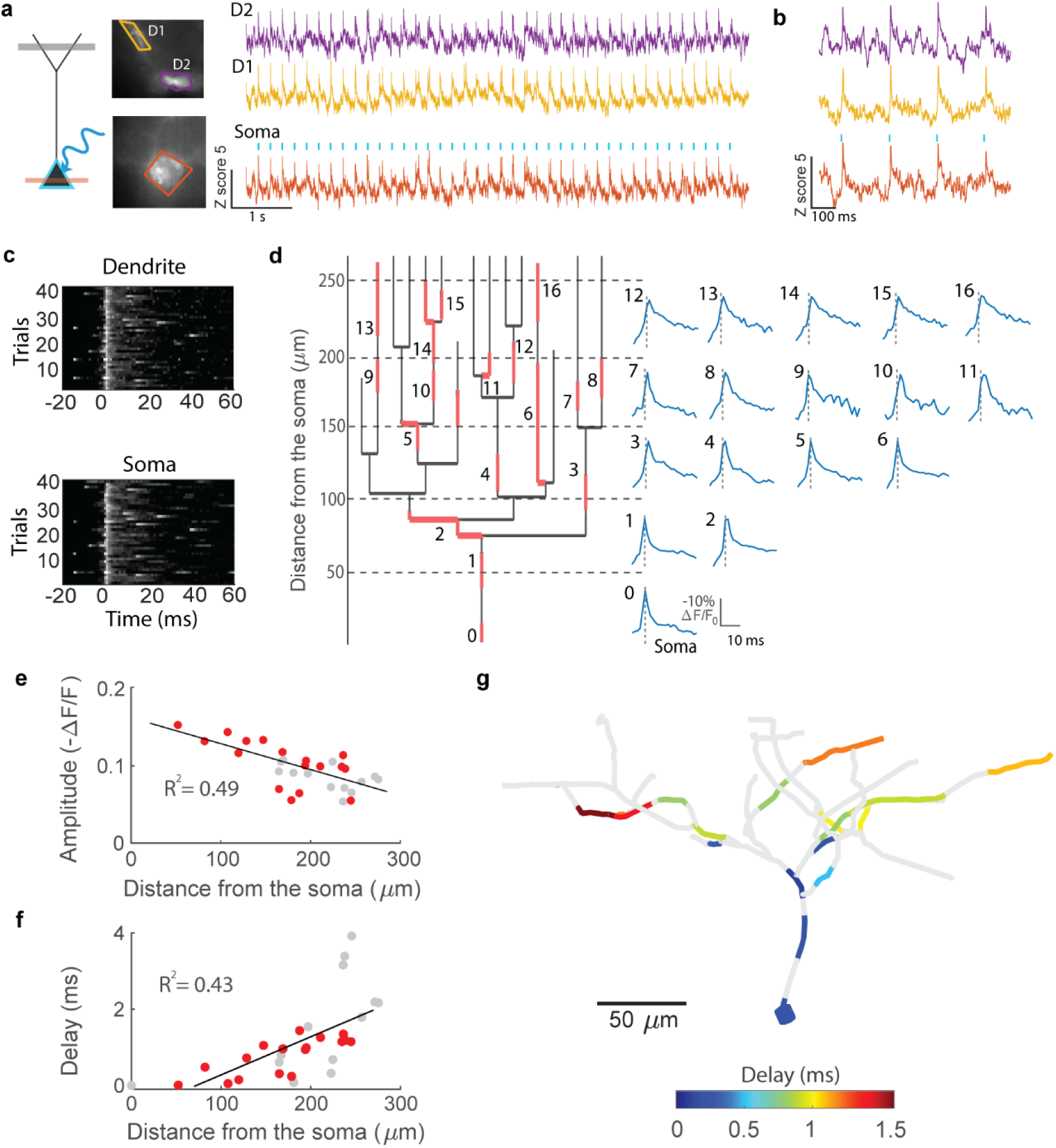
Mapping back-propagating action potentials *in vivo*. **a**, Protocol for measuring bAP delays. Optogenetic stimuli were targeted to evoke somatic spikes and the mean fluorescence responses in the dendrites were calculated, triggered off the soma spike times. Combining many such measurements, always referenced to the soma spike times, yielded a map of bAP delays. **b**, Representative single-trial recordings at the soma and two dendrites. **c**, Stimulus-locked raster illustrates fluorescence dynamics for forty evoked spikes within a single recording. **d**, Dendrogram depicting sixteen dendritic recording sites (red). Right: Spike-triggered averages, triggered off the soma spikes. ΔF/F and time scalebar apply for all STAs. **e**, Amplitude (-ΔF/F) of bAPs vs. contour distance from soma, R^2^ = 0.49 (n = 5 neurons, 5 mice). **f**, Peak delay vs. contour distance from the soma. At distances > 50 μm from the soma, back propagation speed was 0.09 ± 0.02 m/s, R^2^ = 0.43 (n = 30 recording sites, 5 neurons, 5 mice). In (e) and (f), red points from data in (d). **g**, Three-dimensional reconstruction of bAP delays in apical dendrites. See also **Movie S1**.

The bAP peak amplitude decayed to 50% at a contour distance from the soma of 242 ± 30 μm (95% C.I., n = 30 recording sites, 5 neurons, 5 mice, **Fig. 2e**). Back-propagation delays ranged from 1.5 to 4 ms, over contour distances from the soma of 50-280 μm (**Fig. 2f**). Conduction speeds were 0.09 ± 0.02 m/s (95% C.I., n = 30 recording sites, 5 neurons, 5 mice), consistent with prior theoretical predictions ^21^. Finally, we combined multiple recordings to create a comprehensive three-dimensional visualization of bAP propagation throughout a neuron (**Fig. 2g, Movie S1**).

### Dendrite-Dendrite coupling

We next probed the electrical coupling between dendritic branches. Some models of dendritic computation have proposed that dendrites act as distinct computational units ^21,22^. For this to be true, one would expect to observe different voltage dynamics between dendrites of a given cell. We recorded the spontaneous activity in groups of 3 – 5 dendrites simultaneously, with contour-length separations of 70 – 335 μm. In both anesthetized and quietly awake mice, we observed subthreshold fluctuations whose transitions occurred synchronously across dendrites ^23,24^, and whose fine-scale dynamics also appeared correlated between the dendritic branches (**Fig. 3a-c**).

**Fig. 3.**
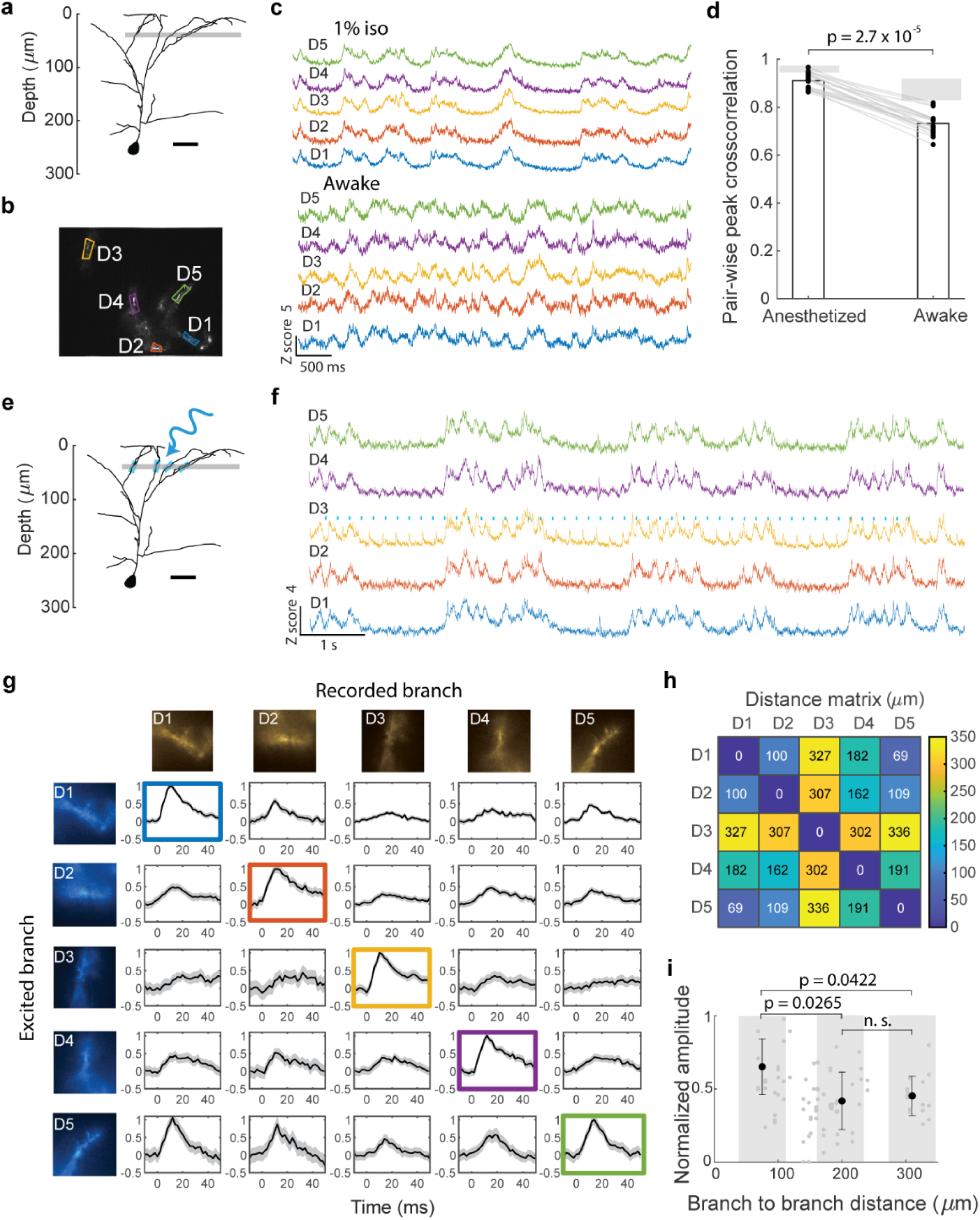
Dendrite-dendrite coupling. **a**, Representative neuron with concurrent recordings from five dendritic branches. **b**, Mean fluorescence of Voltron2 in the recorded dendritic branches. **c**, Recordings of spontaneous activity at (top) 1% isoflurane and (bottom) in the awake state. **d**, Pair-wise cross correlation of recorded branches in anesthetized and awake brain states. Pair-wise cross-correlations were 91 ± 3% (anesthetized, mean ± s.d, n = 23 pairs, 3 neurons, 3 mice) and 73 ± 4% (awake). Shaded bars indicate maximum possible cross-correlation, limited by photon shot-noise. **e**, Blue light targeted to individual apical dendrites evoked small local depolarizations. **f**, Representative traces from a round-robin protocol for estimating dendrite-to-dendrite coupling. Five branches were simultaneously recorded while Dendrite 3 (D3) was stimulated (blue dots). Stimulus-triggered averages of membrane depolarizations provided an estimate of the mean response at each branch. **g**, Matrix showing normalized coupling amplitudes between all branch pairs. Line and shading shows mean ± s.e.m. **h**, Branch-to-branch contour distance matrix of the recording sites, determined from HiLo reconstruction. **i**, Normalized coupling amplitudes at different interbranch distances across the dendritic arbor (n = 92 pairs, 5 neurons, 5 mice). Data points shows mean ± s.d.; p-values: Wilcoxon rank sum test.

To quantify these observations, we calculated pair-wise cross-correlations (**Fig. S8**, Methods) of all branches in the recordings (n = 23 branch-pairs, 3 neurons, 3 mice). Remarkably, dendrites displayed a 91 ± 3% cross-correlation (mean ± s.d.) in the anesthetized state and 73 ± 4% in the awake state (**Fig. 3d**). Photon shot noise degrades the cross-correlation, even if the underlying signals were identical between branches. Accounting for the influence of shot noise (**Fig. S9**, Methods), the upper bounds on the dendrite-to-dendrite cross-correlations were estimated to be 94 ± 2% (anesthetized) and 85 ± 4% (awake). These results imply that the electrical dynamics were largely shared throughout the dendritic tree in L2/3 apical dendrites in both anesthetized and awake states.

The strong correlations between dendrites could be due to strong electrotonic coupling, or to correlated synaptic inputs (or both). To probe dendrite-dendrite coupling directly, we targeted optogenetic stimuli to locally depolarize individual branches, and simultaneously recorded the voltage responses in the stimulated and other non-stimulated branches (**Fig. 3e-f**). Stimulus strength was adjusted to avoid evoking somatic action potentials. We used the eYFP fluorescence from CheRiff-eYFP to confirm that the stimulus light did not spuriously activate non-targeted branches (**Fig. S10**). By averaging the effect of repeated single-branch stimulation (50 repeats, 10 ms pulses, 5 Hz), we calculated a stimulus-triggered average (STA) waveform for the stimulated and non-stimulated branches. This protocol was repeated sequentially across all branches to obtain a dendrite-dendrite coupling matrix (**Fig. 3g**). To estimate the amplitude of each STA, we computed background-robust ΔF/F (Methods) and normalized each row with the amplitude of ΔF/F at the stimulated branch. Inter branch distance was calculated from the three-dimensional reconstructions (**Fig. 3h**).

**Figure 3i** shows dendrite-dendrite coupling amplitude vs. interbranch distance, displaying a 50% decrease after ∼200 μm, and similar ∼50% coupling out to 300 μm. These results imply that most apical dendrites of L2/3 neurons are electrically coupled *in vivo,* allowing for efficient signal transmission within the dendritic tree.

In some trials, stronger dendrite-targeted stimuli evoked spikes. Simultaneous recordings at soma and dendrites in anesthetized mice showed that these spikes usually corresponded to bAPs, although occasionally (2 of 40 trials) dendrite-targeted stimuli evoked dendritic and not somatic spikes (**Fig. S11**).

### Accelerating spike rate favors spontaneous bAP propagation

To study how dendrites mediated back-propagation of action potentials, we first recorded spontaneous activity at the soma and apical dendrites in awake mice. In some cells, we observed spike doublets with a narrow distribution of inter-spike intervals (ISIs, 18 ± 7 ms, mean ± s.d., n = 216 events, 3 neurons, 3 mice, **Fig. 4a**). Examination of individual spike doublets (**Fig. 4b, S12**) as well as the average doublet waveforms (**Fig. 4c**) revealed that the first spike typically failed to propagate to distal dendrites, while the second spike succeeded.

**Fig. 4.**
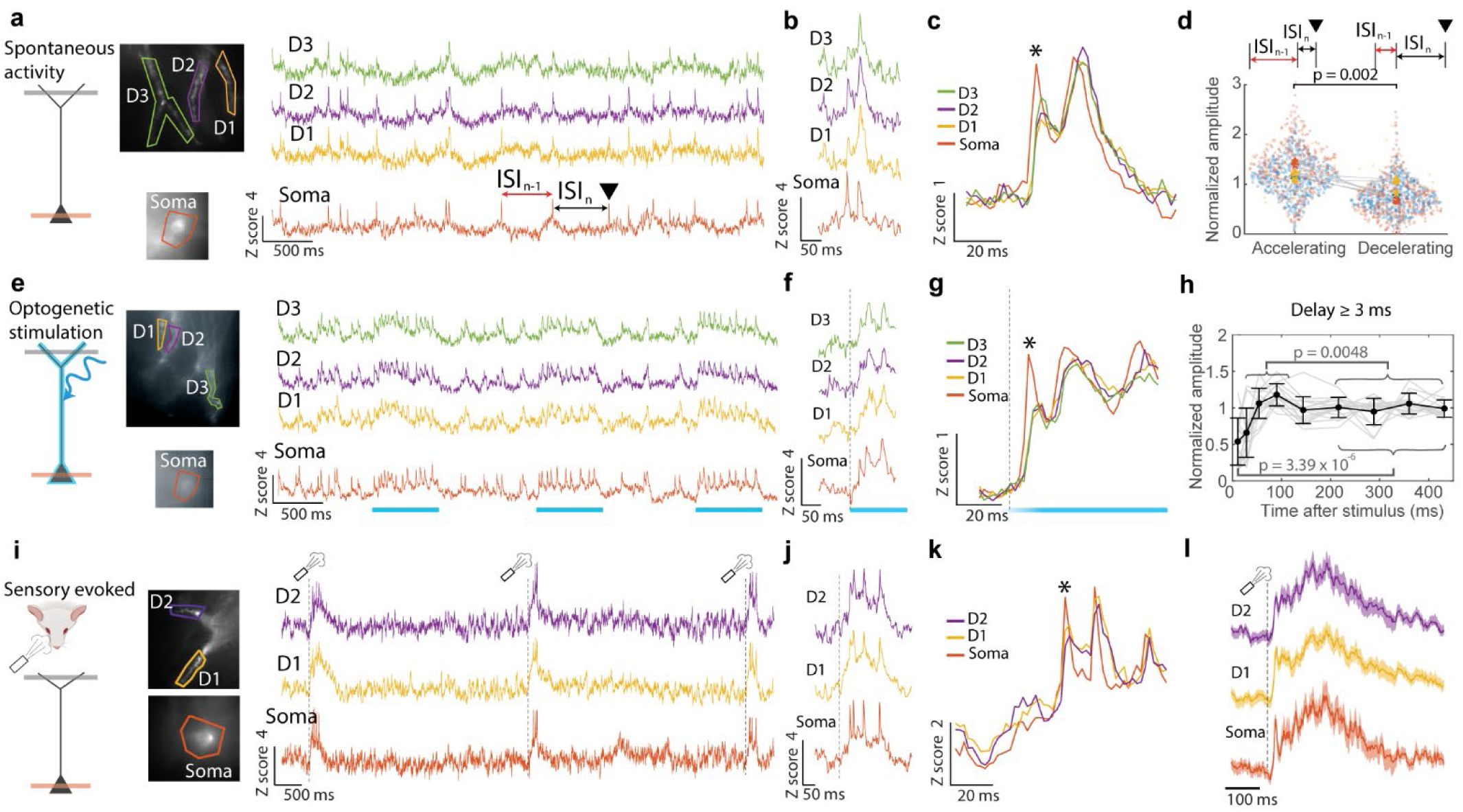
History dependent back-propabation. **a**, Spontaneous activity. Spike timing was characterized by the interspike interval one step (ISI_n_) and two steps (ISI_n-1_) before a reference spike (black triangle). **b**, Single-trial recording of a spike doublet where the first bAP failed to propagate to distal dendrite D3 and the second bAP succeeded. **c**, Spike-triggered average waveform showing attenuation of the first bAP and successful propagation of the second. Events were triggered off the first spike peak at the soma (asterisk, doublet ISIs 18 ± 7 ms, 10 branches, 3 neurons, 3 mice). **d**, Comparison of bAP propagation for accelerating and decelerating spontaneous spike rates. Accelerating: ISI_n-1_ > 75 ms, ISI_n_ < 75 ms; decelerating: ISI_n-1_ < 75 ms, ISI_n_ > 75 ms (n = 186 spike triplets, 3 neurons), p-value: Wilcoxon signed rank test. **e**, Sustained (400 ms) blue light pulses evoked tonic firing at the soma and corresponding bAPs at the dendrites. **f**, Single-trial recording of an optogenetically evoked spike train where the first bAP failed to propagate and subsequent bAPs succeeded. **g**, Spike-triggered average waveform showing attenuation of the first bAP and successful propagation of subsequent bAPs, triggered off the first evoked spike in the soma (asterisk, n = 10 trials, from neuron in **e**). **h**, Biphasic dynamics of bAP amplitude in distal dendrites as a function of time after stimulus onset. Initial bAPs were suppressed, followed by a window of enhanced bAP amplitude, and then a decrease (n = 16 trials, 7 neurons, 7 mice). P-values: Wilcoxon signed rank test. **i**, Air puff sensory stimulation evoked bursts of spikes in the soma and bAPs in the dendrites. **j**, Single-trial recording of a sensory-evoked burst where the first bAP failed to propagate and subsequent bAPs succeeded. **k**, Spike-triggered average waveform showing attenuation of the first bAP and successful propagation of subsequent bAPs in distal dendrites (n = 13 trials). Events were triggered off the first spike peak at the soma. **l**, Stimulus-triggered average responses showing a fast depolarization followed by a sustained plateau potential. Line and shading: mean ± s.d.

To quantify this effect, we classified changes in spike rate in trios of consecutive spikes by the ISI one step (ISI_n_) and two steps (ISI_n-1_) before a reference spike (arrow in **Fig. 4a**). We defined accelerating trios as ISI_n-1_ > 75 ms and ISI_n_ < 75 ms and *vice versa* for decelerating (**Fig. 4d**). The third bAP in accelerating trios experienced amplification relative to the second bAP, while the third bAP in decelerating trios experienced suppression relative to the second (10 branches, 3 neurons, 3 mice). Thus, spontaneous bAP propagation *in vivo* is influenced by the rate of change of firing, favored by an accelerating spike rate.

### Accelerating spike rate favors optogenetically triggered bAP propagation

Since spontaneous spikes are evoked by synaptic inputs, the observations above could reflect either intrinsic properties of dendritic excitability, or an aspect of the synaptic inputs around the time of the spike doublet (e.g. changes in shunting from balanced excitation and inhibition). To resolve this ambiguity and to gain independent control of the spike timing, we stimulated the soma with sustained (> 400 ms) pulses of blue light while recording from the soma and dendrites (**Fig. 4e**). These pulses produced tonic firing at the soma and corresponding bAPs in the dendrites.

We calculated the average bAP amplitudes in the dendrites as a function of time after stimulus onset. As in the spontaneous activity, the first bAP in each spike train was highly attenuated in the dendrites relative to the soma (**Fig. 4f-g**). There was a window from 50 –150 ms after stimulus onset where bAP amplitude increased; then bAP amplitude decreased again (**Fig. 4h**). This biphasic modulation of bAP amplitude was larger in dendrites further from the soma (as measured by mean propagation delay) than in more proximal dendrites (**Fig. S13**, 25 branches, 7 cells, 7 mice). These results establish that the history-dependent modulation of bAP propagation is an intrinsic feature of dendritic excitability.

### Accelerating spike rate favors sensory evoked bAP propagation

Finally, we studied whether the propagation patterns observed for spontaneous and optogenetically evoked spikes also applied to sensory evoked activity. Air puff stimulation to the contralateral whiskers in awake mice evoked bursts of activity in both somas and dendrites (**Fig. 4i**) with all detected dendritic spikes initiating as bAPs from the soma. Examination of the single-trial responses (**Fig. 4j**) as well as the average evoked waveforms (**Fig. 4k**) revealed that the first spike in the burst typically failed to propagate to distal dendrites, while subsequent spikes succeeded (n = 13 trials). The average dendritic waveform showed a fast sensory-evoked upstroke, followed by a sustained plateau depolarization which lasted ∼400 ms, substantially longer than the stimulus. This response motif suggests NMDA receptor engagement ^9^, an important mediator of dendritic excitability which is not engaged by direct optogenetic stimulation.

## Discussion

Techniques to study subcellular voltage dynamics *in vivo* are critical for understanding neural computation and plasticity. Here we presented an all-optical approach to investigate dendritic signal processing, focusing on Layer 2/3 pyramidal neurons in the barrel cortex of awake and anesthetized mice. We combined dual-plane voltage imaging with targeted optogenetic stimulation to study bidirectional electrical couplings between dendrites and soma, and between dendrites and dendrites, at both the subthreshold and suprathreshold level. Overall, our findings indicate that electrical dynamics are largely shared throughout the dendritic tree. In our experiments, the primary role of dendritic nonlinearities appeared to be in history dependent modulation of bAP propagation, rather than in dendritic integration.

The biphasic bAP propagation efficiency in response to a step-increase in spike rate (**Fig. 4h**) is similar to the dynamics observed in an analogous experiment in CA1 pyramidal cells in acute slices ^25^. In that work, pharmacological perturbations ascribed the initial increase in bAP propagation to A-type potassium channel inactivation ^26^, and the subsequent decrease in bAP propagation to slower sodium channel inactivation ^27^. The combined biphasic bAPs transmission function suggests that the dendrites act as a spike-rate accelerometer: neither isolated spikes nor sustained spike trains are transmitted efficiently, but during an increase in spike rate there is a transient window for efficient bAP propagation. This motif has important implications for learning rules: it suggests that bursts, not individual spikes, will be most effective at engaging plasticity ^28^, and provides a means for sensing causal influences of synaptic inputs on spike rate, regardless of basal spike frequency. We speculate that the loss of bAP propagation efficiency during sustained high-frequency spiking may implement a feedback to prevent runaway plasticity.

Our observation of broadly distributed and highly correlated subthreshold voltages may be context specific. For example, dendritic length constants may be shortened in animals engaged in specific tasks or in altered states of arousal. Furthermore, other neuron types, such as L5 or CA1 pyramidal cells, have more extensive dendritic arbors which may support localized electrical signaling. Finally, the nonlinear relationships between voltage and calcium permit localized Ca^2+^ signals even in the presence of broadly distributed voltage. We recently observed that subtle changes in bAP amplitude could drive dramatic branch-to-branch differences in Ca^2+^ in L2/3 pyramidal cells in acute slices, a consequence of nonlinear activation of voltage-gated Ca^2+^ channels ^29^. Similarly, activation of NMDA receptors could mediate local Ca^2+^ accumulation despite broadly distributed voltage dynamics.

A deeper understanding of electrical propagation in dendrites could help elucidate the relations between dendritic biophysics and information processing. Critical next steps will be to combine imaging of voltage with recordings of neurotransmitters, neuromodulators, Ca^2+^ dynamics, and signaling pathways in animals engaged in tasks which activate diverse brain states.

## Acknowledgements

We thank A. Preecha and S. Begum for technical assistance and B. Sabatini for helpful discussions. This work was supported by a Vannevar Bush Faculty Fellowship, a Brain Research Foundation Scientific Innovations Award, the Harvard Brain Science Initiative, and NIH grants R01-NS126043 and R01-MH117042. J.D.W.-C. is a Merck Awardee of the Life Sciences Research Foundation. J. B. G., S. E. P. and L. L. are supported by the Howard Hughes Medical Institute. J.D.W.-C. acknowledges Dalia P. Ornelas-Huerta for useful discussions.

## Methods

### DNA constructs

The Cre-dependent Optopatch construct was generated using standard molecular cloning techniques. All the new constructs and their sequences are available on Addgene.

### Animals

All procedures involving animals were in accordance with the National Institutes of Health Guide for the Care and Use of Laboratory Animals and were approved by the Harvard University Institutional Animal Care and Use Committee (IACUC). The *in utero* electroporation (IUE) surgery was made as described previously ^30^. Briefly, embryonic day 15.5 (E15.5) timed-pregnant female CD1 mice (Charles River, Wilmington, MA) were deeply anesthetized and maintained with 2% isoflurane. The animal’s body temperature was maintained at 37 °C. Uterine horns were carefully exposed, and periodically rinsed with warm sterile phosphate-buffered saline (PBS). The plasmid DNA was diluted with PBS (2 μg/μL; 0.005% fast green), and 1 µL of the mixture was injected into the left lateral ventricle of pups. Electrical pulses (40 V, 50 ms duration) were delivered five times at 1 Hz using a tweezers electroporation electrode (CUY650P5; Nepa Gene, Ichikawa, Japan). Injected embryos were placed back into the abdominal cavity, and the surgical wound was sutured with PGCL25 absorbable sutures (Patterson, Saint Paul, MN).

### Cranial window surgery

Surgeries were conducted on 4-week-old mice of both sexes, following the protocol outlined by ^31^. The surgical procedure began by exposing the skull, followed by a 3 mm circular craniotomy at coordinates 3.3 – 3.4 mm lateral and 1.6 mm caudal relative to the bregma. The craniotomy was made using a biopsy punch (Miltex). The dura mater was then carefully removed while taking measures to prevent capillary bleeding. Following this, a 3 mm round cover glass (Thomas Scientific, 1217N66), pre-glued to a custom-made stainless-steel adapter (Tecan, **Fig S3**), was inserted into the opening. This setup aimed to reduce the thickness of the glass at the brain-objective interface. To prevent any relative movement between the brain and the adapter, a gentle pressure was applied, and the outer ring of the adapter was securely glued. All subsequent experimental procedures were carried out at least one-week post-surgery, ensuring that the health of each mouse was stable and had been duly confirmed.

### Optical system

Experiments were performed on a custom-built upright fluorescence microscope (**Fig. S1**). Illumination was delivered by lasers centered at 488 nm (Coherent, OBIS, 60 mW) and 594 nm (Cobolt Mambo, 100 mW). Laser light was combined using a dichroic mirror (IDEX, FF506-Di03-25×36) and sent through an acousto-optic tunable filter (Gooch and Housego, TF525-250-6-3-GH18A) for modulation of each wavelength. The beams from each color were then separated by another dichroic mirror (IDEX, FF506-Di03-25×36) and sent to two digital micromirror devices (DMD, Vialux, V-7000). This configuration allows dynamic stimulation and recording of independent areas at two wavelengths. Light from both DMDs was merged using a dichroic mirror and converged at an intermediate image plane, which served as a point for validating the optical alignment. An excitation tube lens (Thorlabs, LA1417-A) collected light from this intermediate image plane and sent it to a 25× water immersion objective, 1.05 numerical aperture (Olympus XLPLN25XWMP2 25×, 1). Fluorescence was collected by the same objective and separated from excitation light via a multi-band dichroic mirror (IDEX, Di01-R405/488/594-25×36, three bandpasses). The dichroic was carefully mounted to minimize mechanical stress which could induce astigmatism in the image.

A 50-50 beamsplitter (Thorlabs, BSW26R) split the light into two image paths, each of which was passed through an emission filter (Chroma, ET645/75m, bandpass) placed after the camera, a tube lens (Thorlabs, TTL200-A) on an axial translation stage, and imaged onto a scientific CMOS camera (Hamamatsu, ORCA-Fusion and ORCA-Flash). The cameras were synchronized by frame-trigger pulses from a data acquisition system (National instruments, NI-PCIe-6343) referenced to the output clock signal of the camera. This same clocking synchronizes the experimental waveforms that control the AOTFs and shutters. Field of view registration between the DMDs and cameras were performed by an affine transformation between a calibration pattern projected on to the DMDs consisting of three points.

Overall focus was achieved by moving the sample axially relative to the objective, while offset in the focal planes was achieved by axial translation of the tube lens in front of the second camera. A kinematic mirror before the tube lens of the second camera allowed lateral offset between the two camera fields of view. Our scheme allows arbitrary control of the field of view of the second camera across 400×400×300 μm^3^ (**Fig. S14**).

## Experimental protocols

### Cranial window adapter

Commonly used three-layer cranial windows ^31^, consisting of three coverslips glued together, are outside the thickness range of typical microscope objective correction collars, leading to spherical aberration. To mitigate this source of aberration, stainless steel adapters (Tecan) were designed to mount a single #1 coverslip against the brain surface. The adapter consists of a washer-like structure with a 150 μm protrusion in an inner ring, where the #1 coverslip is glued (**Fig. S3**). Additionally, to counteract spherical aberration in dendrites, we incorporated a technique of imaging a dendrite and adjusting the focus in a scanning motion. Adjustments to the correction collar altered the relative symmetry of the image between the in-focus and out-of-focus planes typically displayed as a different blurred image. Correctly adjusted configurations display similar image shapes at both out-of-focus planes.

### Animal tip-tilt

Coma aberration associated with sample tilts were compensated by adding a goniometer stage (Optics Focus, MAGXY-60L-4-6M) under the mouse platform. Coma aberration in dendrites is displayed as sideways blurring of the fluorescence signal. We adjusted the goniometer by observing the side blurring and refocusing for every tilt step. A correctly adjusted tip-tilt minimizes sideways blurring along the tilted axis.

### Blood vessel identification

Neurites and somas below a blood vessel are subject to local variations in the absorption and scattering of the stimulating and fluorescent light, resulting in stochastic signal variations ^32^. Blood vessels were visualized by injecting a fluorescent tracer (Sigma, 74817, FITC-CM-Dextran) and avoided in our recordings. We retro-orbitally delivered 100 μL of solution at a concentration of 50 mg/mL of the dye. Excitation at 488 nm allowed the tracking of the capillaries. Additionally, two photon microscopy z-stacks could be used to localize all capillaries within a volume.

### Dye injection and depth penetration

We characterized the light penetration across the wavelengths we used in our experiments. For the blue spectrum, we retro-orbitally injected the FITC-Dextran dye and recorded a 1P z-stack. Capillaries deeper than 50 μm were not resolvable. For the red side of the spectrum, we retro-orbitally injected 100 μl of a red dye (ThermoFisher, D22914, Alexa Fluor^TM^ 647-Dextran) at 50 mg/ml and recorded a 1P z-stack. Imaging with red shifted wavelengths allowed imaging of capillaries down to 300 μm of brain tissue (**Fig. S7**)

### One-photon light targeting with digital micromirror devices (DMD)

We employed optogenetic activation with blue light to selectively target either apical dendrites or cell bodies. Optical stimulation of individual dendritic branches required up to ∼100-fold greater blue light intensity compared to stimulating the soma. For targeted soma activation, we selected regions devoid of apical dendrites to prevent spurious dendritic stimulation. For targeted dendrite stimulation, we restricted the blue light to just the targeted branch, to minimize spurious activation of the soma or other branches by out-of-focus or scattered light. For voltage recordings, the orange light was meticulously confined to the recording sites, which led to a decrease in background fluorescence.

### Imaging sessions

Imaging experiments started 3-4 weeks after the cranial window surgery. JFX_608_-HaloTag ligand ^33^ (100 μl, 1 mM) was retro-orbitally delivered 24 hrs before the imaging session. We chose a red shifted dye due to its compatibility with our blue shifted Channel-rhodopsin, voltage sensitivity and deep tissue penetration (**Fig.S7)**. Neurons remained labeled for up to one week. Injection of a lower volume (30 μl and 50 μl) did not substantially affect the measurement of voltage activity. For imaging the voltage indicator, we used a 594 nm laser (Cobolt Mambo, 100 mW). Illumination intensity was 25 mW/mm^2^ at the sample plane.

### Cell health validation

Cell health was assessed by the response of a cell to blue light steps of increasing intensity. This protocol activated the neuron and led to spiking rates correlated to the light intensity. We performed this validation before and after each imaging session.

### Imaging anaesthetized animals

Imaging typically started 2-3 weeks after surgery. Mice were lightly anaesthetized (0.7–1% isoflurane) and head-fixed under the upright microscope using a titanium head plate. Eyes were kept moist using ophthalmic eye ointment. The body temperature was continuously monitored and maintained at 37 °C using a heating pad (WPI). A typical imaging session lasted 2–3 h, after which the animals quickly recovered and were returned to their home cage.

### Imaging awake animals

Head-fixed animals were imaged while awake on a custom-built motorized treadmill. Before imaging sessions started, mice were habituated to head restraint by training them every 24 h, for 15–30 min. During each training session mice were first habituated to head restraint until completely relaxed. Whisker stimulation was applied to the contralateral whiskers using an air puff of 100 ms and 45 psi (Picosprizer III, Parker). Stimuli were performed during imaging of the somatic and dendritic compartments at 4 second intervals.

## Data analysis

### Three-dimensional motion correction

Transverse xy motion correction was performed by using a MATLAB implementation of NoRMCorre (https://github.com/flatironinstitute/NoRMCorre). For z-motion correction, before each imaging session a z-stack was acquired with axial displacements up to ±10 μm from the focal plane. After a voltage imaging acquisition, each frame of the recording was compared to the calibration z-stack to determine the axial displacement. Frames that deviated by more than 3 μm in axial displacement were excluded from analysis.

### Fluorescence signal extraction

Videos were corrected for photobleaching by dividing the video by an exponential fit of the mean fluorescence. Activity-based image segmentation was performed on the spiking component of the signal. We applied PCA to a region of interest around each individual branch to isolate the voltage signal from residual motion or blood-flow artifacts. Regions of the movie with blood vessels were masked.

### One-photon HiLo for morphology reconstruction

Three-dimensional mapping of neurons was performed using HiLo ^20^ at a succession of z planes. The three-dimensional structural data cube was imported to Neutube and semiautomatic reconstruction was performed. In some instances, we also performed 2P structural microscopy. We did not observe significant differences between our 1P and 2P mapping approaches. Basal dendrites were typically not resolved in our recordings.

### Robust estimate of ΔF/F

For analysis of bAP amplitude of **Fig. 2** and the small local depolarization events in **Fig. 3**, we created a spike-triggered average (STA) movie comprising the average of all depolarization events. An initial estimate of the STA waveform for each dendritic branch was calculated by averaging the pixel values in a region of interest comprising 5% of pixels with greatest mean brightness. Maps of the baseline (F) and spike-dependent (ΔF) signals were then calculated pixel-by-pixel by linear regression of the STA movie against a normalized copy of the initial STA waveform estimate. For the region surrounding pixels with the highest ΔF, a pixel-by-pixel plot of F vs. ΔF was fit to a line via linear regression. The inverse of the slope was the calculated ΔF/F.

### Contribution of shot noise to imperfect dendrite-to-dendrite cross-correlation

To estimate the contribution of photon shot-noise to the less-than-perfect correlation, we took the region of interest around each dendritic branch and split it into two signals derived from interleaved checkerboard patterns of pixels (**Fig. S9**). These two signals are expected to be nearly identical, except for independent realizations of the photon shot-noise. The cross-correlation of these reference signals set an upper bound on the expected cross-correlation between signals from distinct dendritic branches.

### Dendrite-dendrite coupling matrix

Round robin stimulation of individual branches with pulsed optogenetic stimulus was used to compute a coupling matrix between branches. By calculating the STA of the stimulated and recorded branches we obtain the rows of this matrix. We obtain the voltage response of the stimulus with our estimates of ΔF/F. To control for variation in CheRiff expression and optogenetic stimulus strength, we normalized the amplitudes of all recordings (rows) by the amplitude of the response at the corresponding stimulated branch. Hence the amplitude of the diagonal elements of this coupling matrix is identically 1.

### Analysis of bAP failures during spontaneous activity

We first found each spike at the soma, and the corresponding events in each dendritic recording. For each dendritic compartment, we normalized bAPs by their mean amplitude, to allow analysis of variations in bAP amplitude within and between recording sites. Inter-spike intervals were calculated at the soma and classified into “accelerating” or “decelerating” triplets. The amplitude of the third spike in each triplet was determined and was classified based on whether the preceding two ISIs were accelerating or decelerating.

### Analysis of bAP failures during optogenetic stimulation

We first found all optogenetically evoked spikes at the soma and identified the corresponding events in the dendritic recordings. For each dendrite, bAP amplitdues were normalized by the mean bAP amplitude in the final half (200 – 400 ms) of the stimulus period, a time when we observed that bAP amplitudes were stable. Spikes and their corresponding bAPs were sorted into time bins relative to the optogenetic stimulus onset. Within each time-bin, we calculated a spike-triggered average waveform at the soma and the dendrites, triggered off the spike times at the soma, and then calculated the mean amplitude of the bAP in each dendritic compartment. Dendritic compartments were assorted into groups based on the time delay between the peak of the soma spike and the peak of the local bAP. Mean bAP amplitudes were calculated for electrotonically proximal dendrites (conduction delay < 3 ms) and for electrotonically distal dendrites (conduction delay > 3 ms).

## Supplementary figures

**Figure S1.**
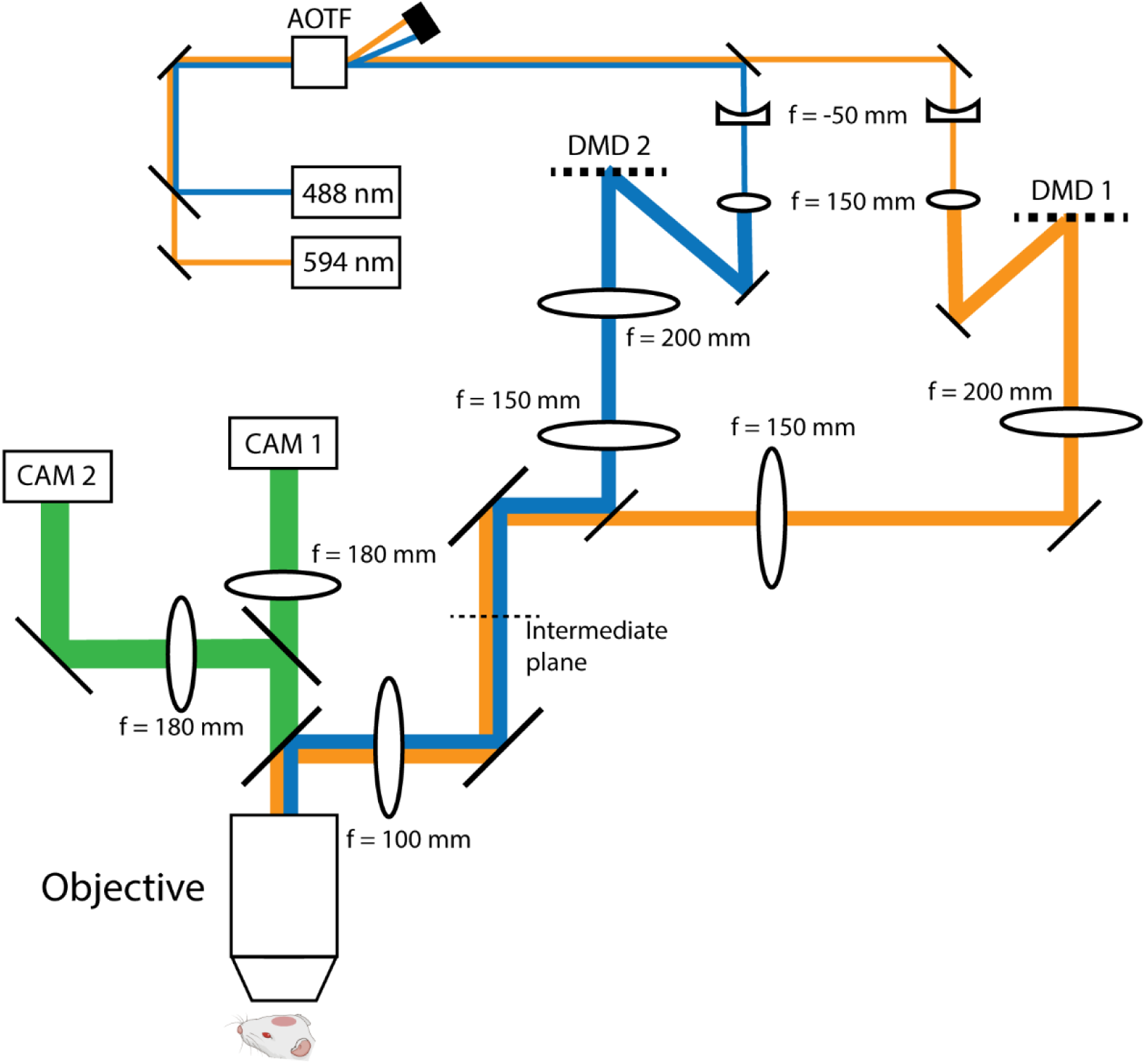
Optical layout. 488 and 594 nm light is amplitude modulated by an acousto-optic tunable filter (AOTF) and expanded by a 1:3 telescope. Blue and orange light illuminate two separate digital micromirror devices (DMDs). The image of each DMD is reimaged at an intermediate plane and relayed to the sample. Fluorescence (green) is redirected by a 50:50 beam splitter to two cameras. Adjustment of the tube lens (f=180 mm) allows control of the imaging field of view of each camera.

**Figure S2.**
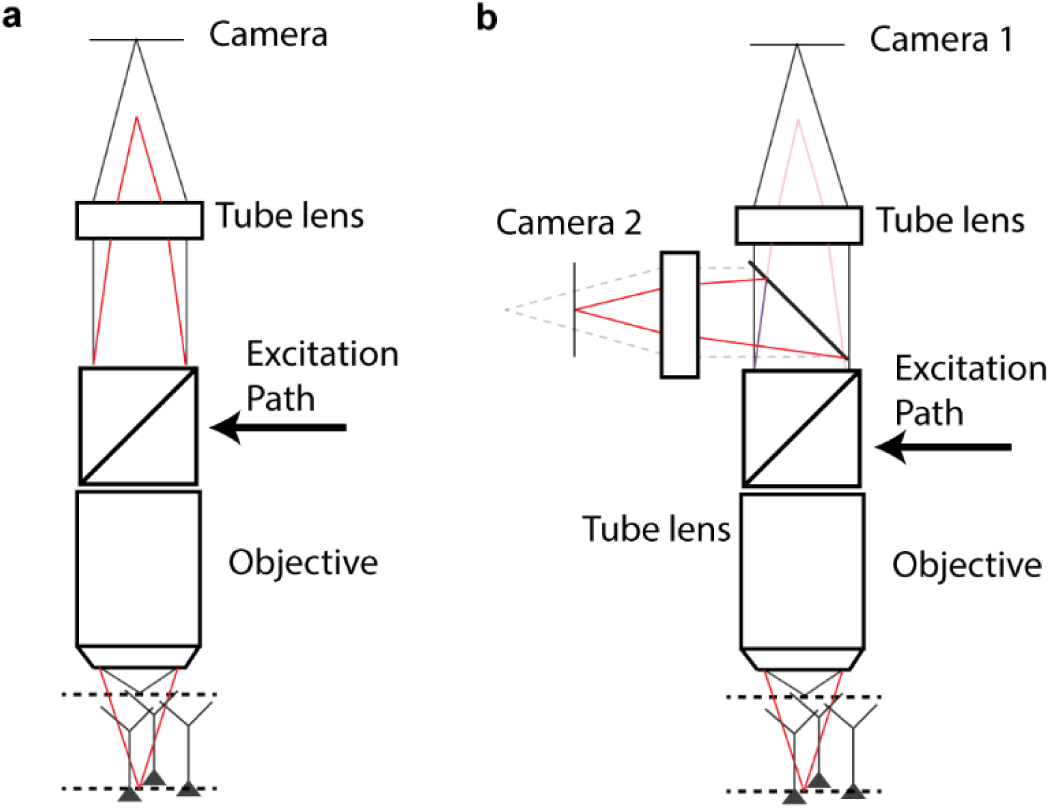
Simultaneous 1P imaging of two focal planes. **a**, Light from the soma couples to the objective and is reimaged at a different focal plane, located closer to the objective (Red). **b**, By adding a beam splitter, we image this plane to a second camera, allowing simultaneous recordings of somas and dendrites.

**Figure S3.**
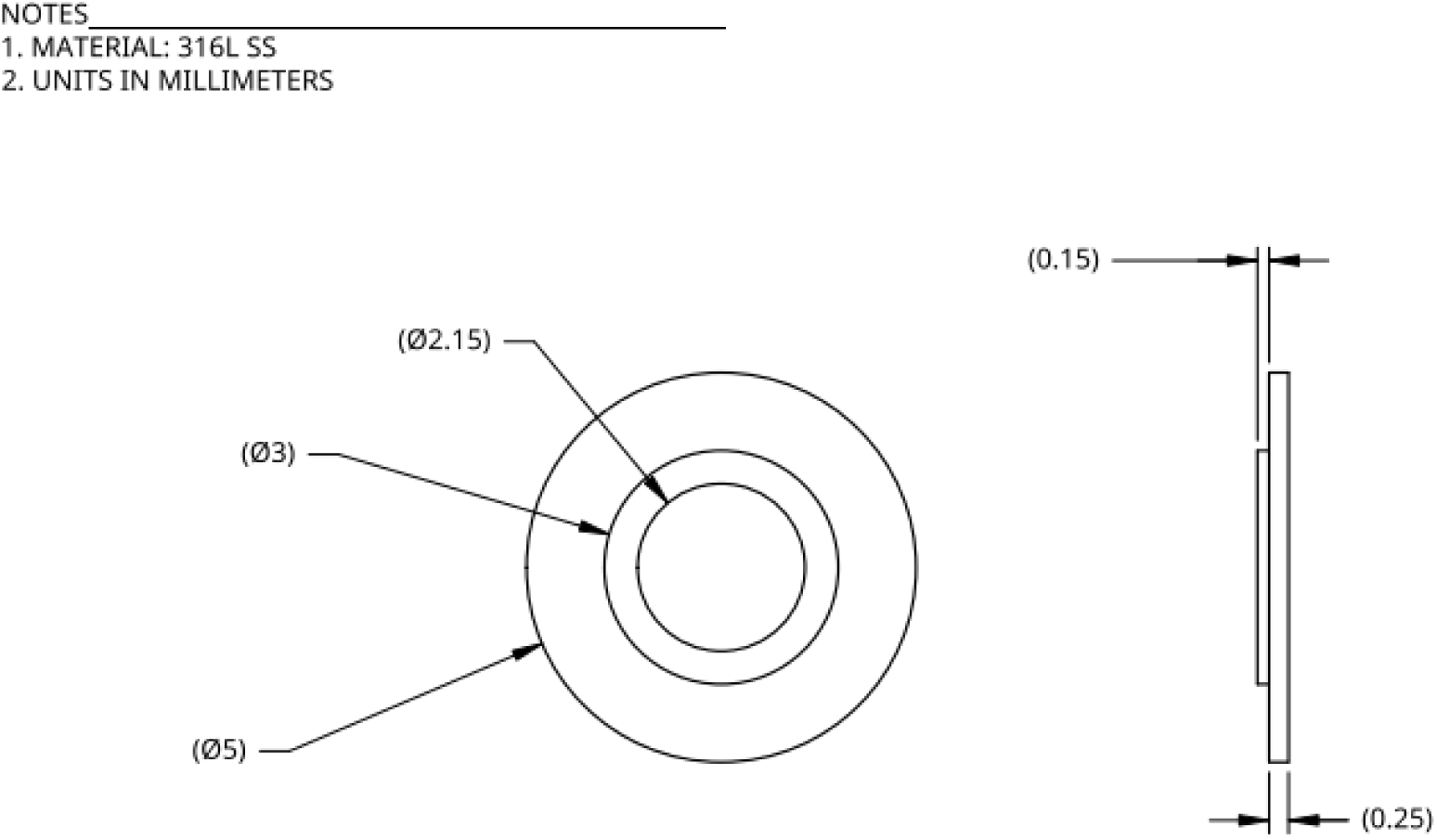
Stainless steel adapter. A cover slip of 3 mm diameter and 150 µm thickness is glued to the inner ring of 3-2.15 mm diameter. This adapter minimizes the spherical aberration from the brain-glass-water interface. All units are in millimeters.

**Figure S4.**
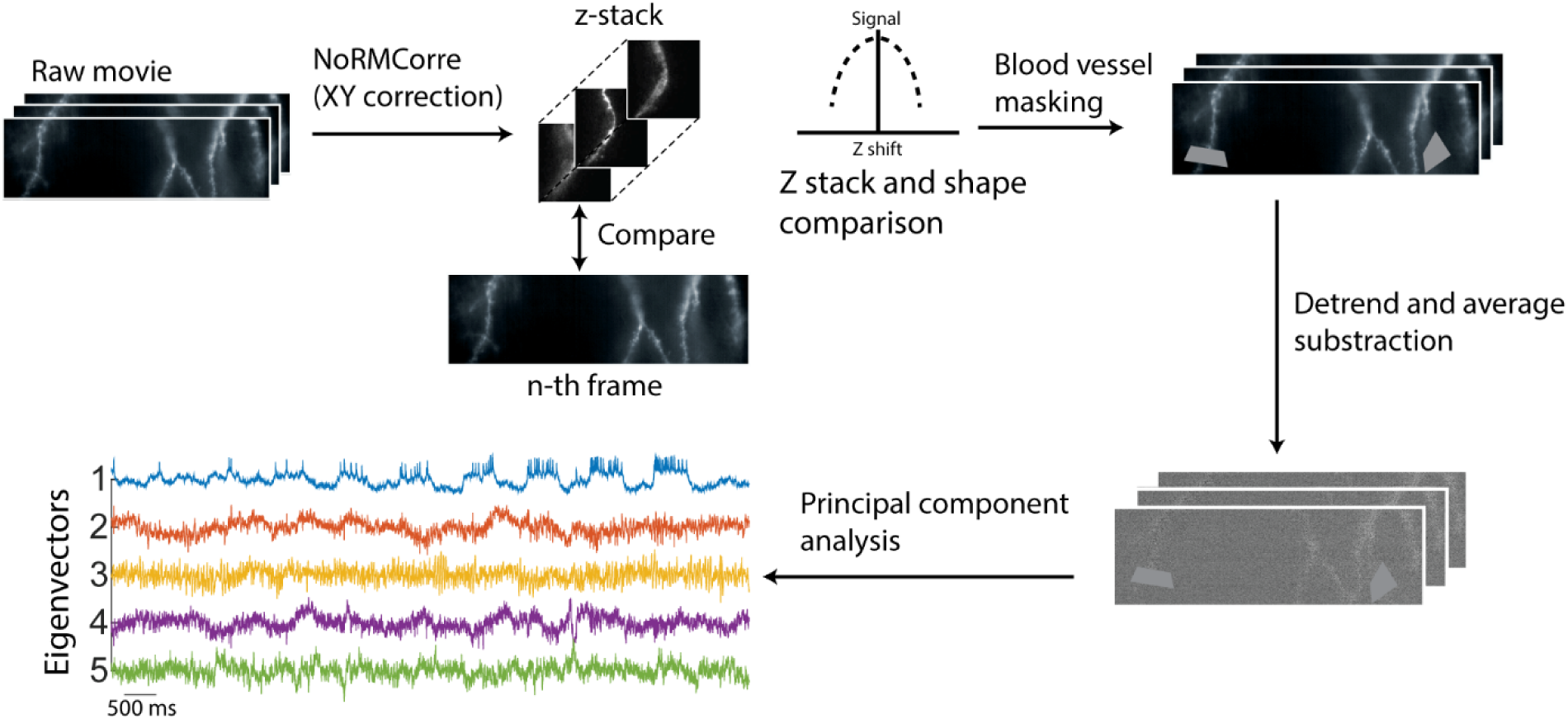
Analysis pipeline. Raw movies are motion corrected for transverse (XY) and axial (Z) drifts. Blood vessels are masked, and the movie is detrended. The average-subtracted movie is processed with Principal Component Analysis (PCA)and the main amplitude component is kept. This operation is repeated on a per-branch basis.

**Figure S5.**
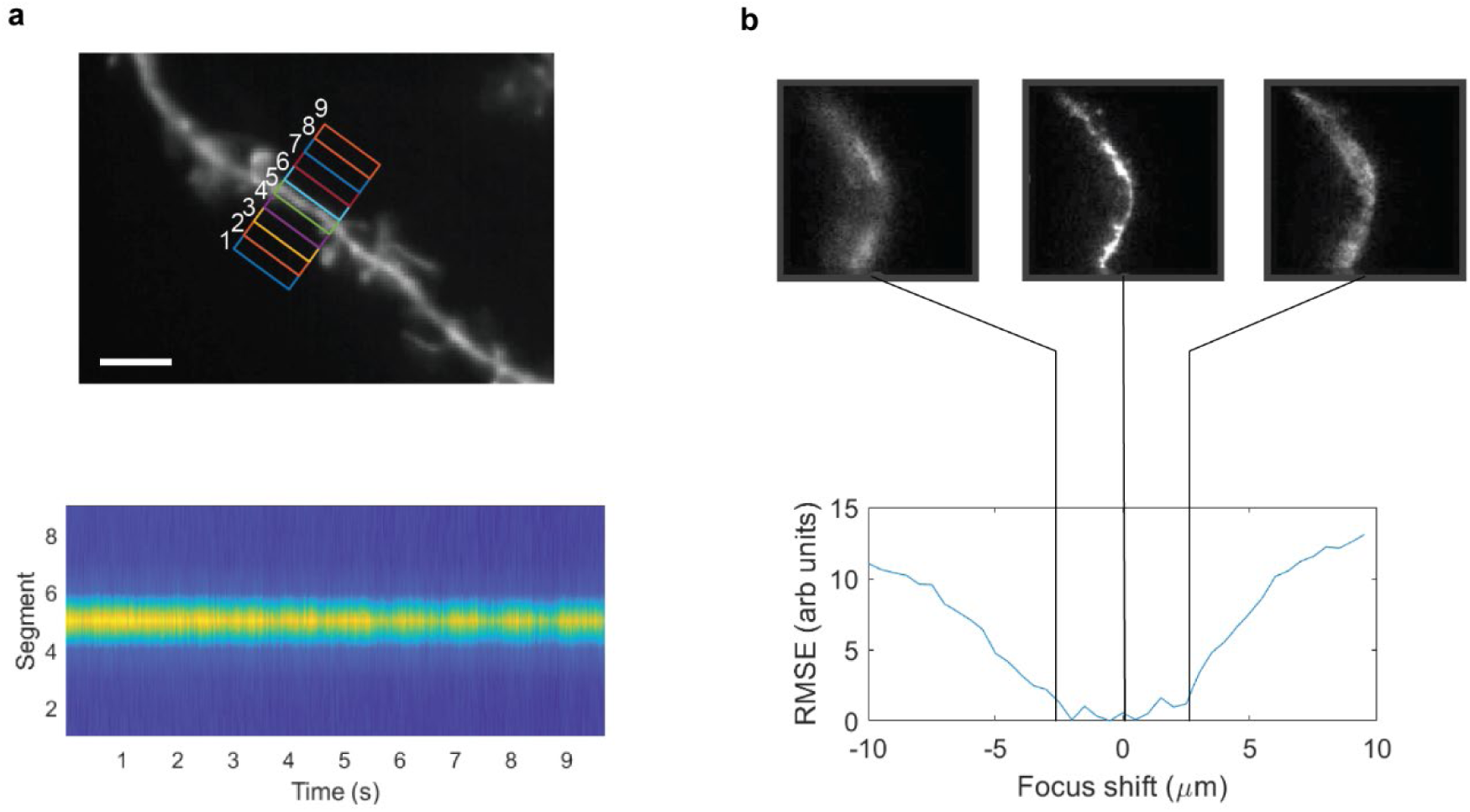
Signal localization in one-photon structured illumination microscopy *in vivo*. We subdivide a region transverse to a dendrite into 9 segments of 1.2 μm width and measure the fluorescence vs. time of each segment. The signal is spatially localized to 2.4 μm in the transverse direction. Scale bar of 5 μm. **b**, For axial motion correction, we first acquire images of the dendrite as a function of axial defocus and compute the root mean square error (RMSE) of each image relative to the in-focus image. The signal is maintained to ±2.5 μm defocus. We compare each frame of our raw movie with the Z stack calibration. Any frames with axial shift that exceed this ±2.5 μm shift are excluded from analysis.

**Figure S6.**
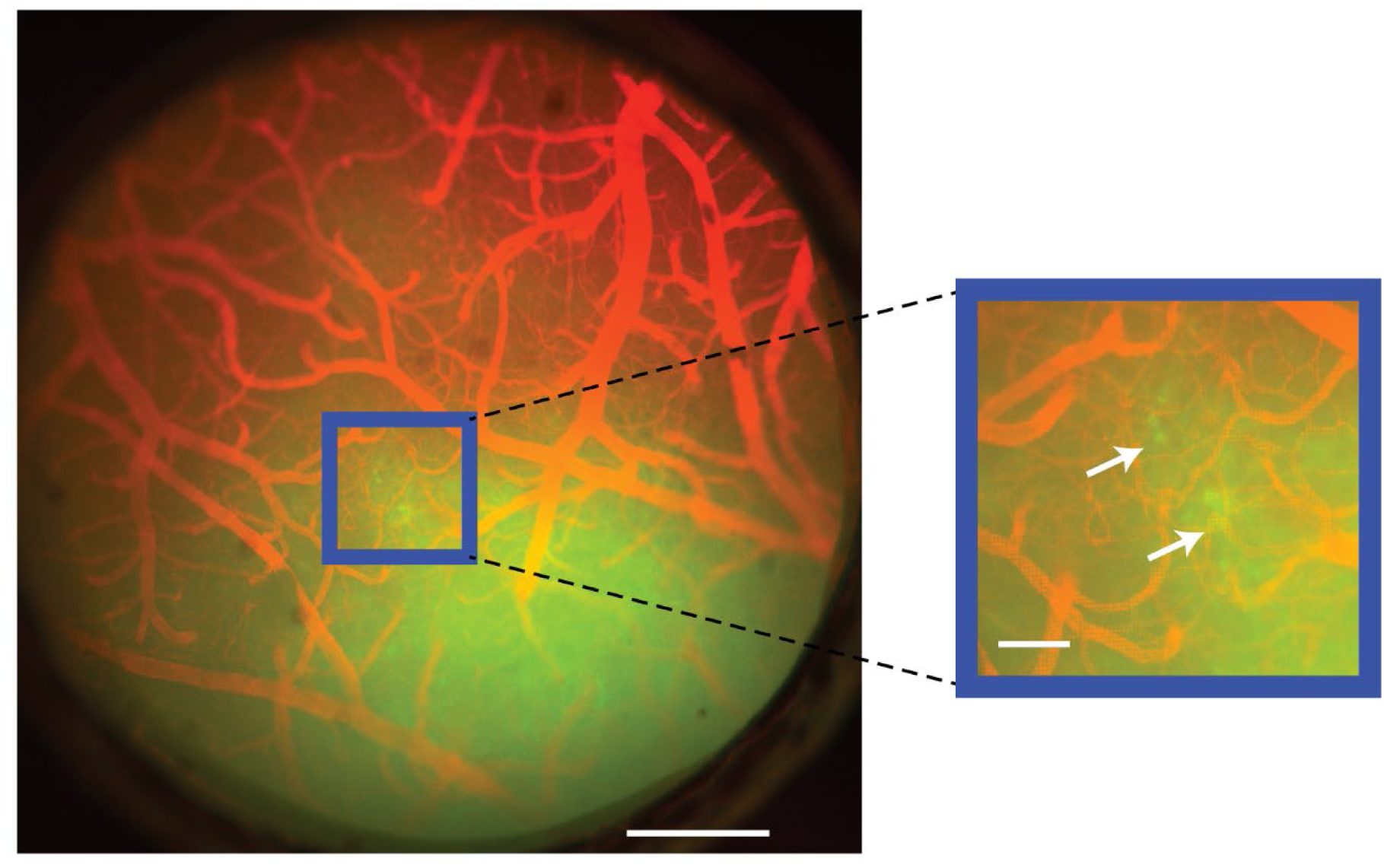
Blood vessel tracking. Blood vessel visualization was performed by injecting a fluorescent tracer. Scale bar 500 μm. We avoided recordings close to capillaries due to increased noise caused by scattering of flowing blood cells. Red: blood vessels, green: voltage indicator. Inset: Zoom in and arrows showing somas. Scale bar 250 μm.

**Figure S7.**
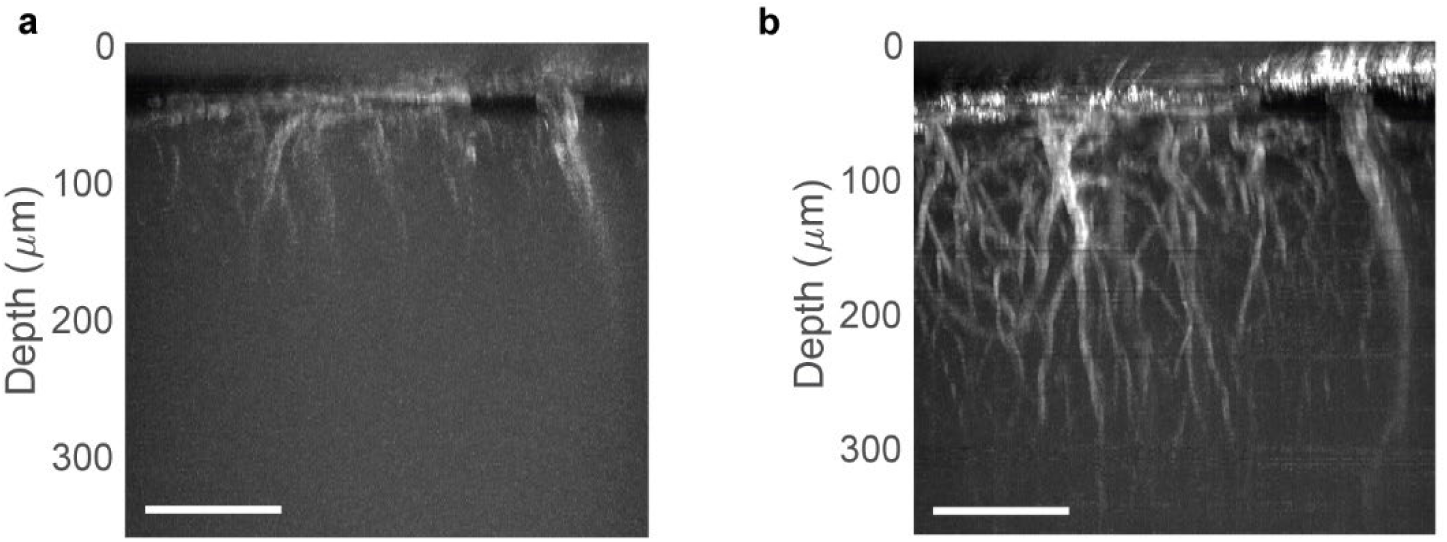
Wavelength-dependent penetration depth of one-photon structured illumination imaging. Side projection of one-photon z-stacks of labeled blood vessels acquired via HiLo. Images show the same sample and field of view, labeled with two different dyes. **a**, Imaging at 488 nm and **b**, Imaging at 635 nm. Scale bar 100 μm.

**Figure S8.**
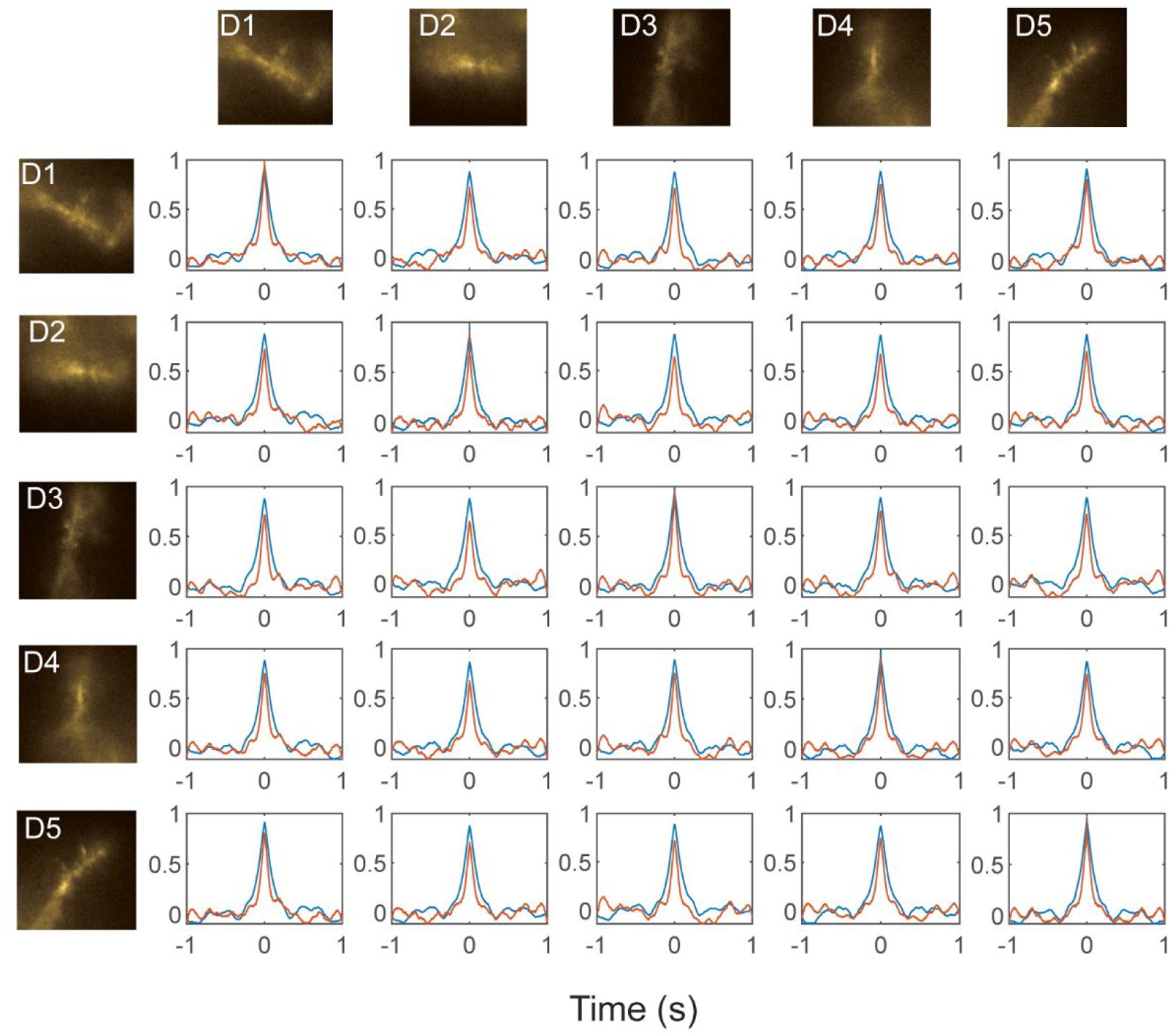
Cross-correlation matric of spontaneous activity. Pair-wise cross correlation matrix of the neuron in Fig. 3. Blue: anesthetized (1% isoflurane); Orange: awake.

**Figure S9.**
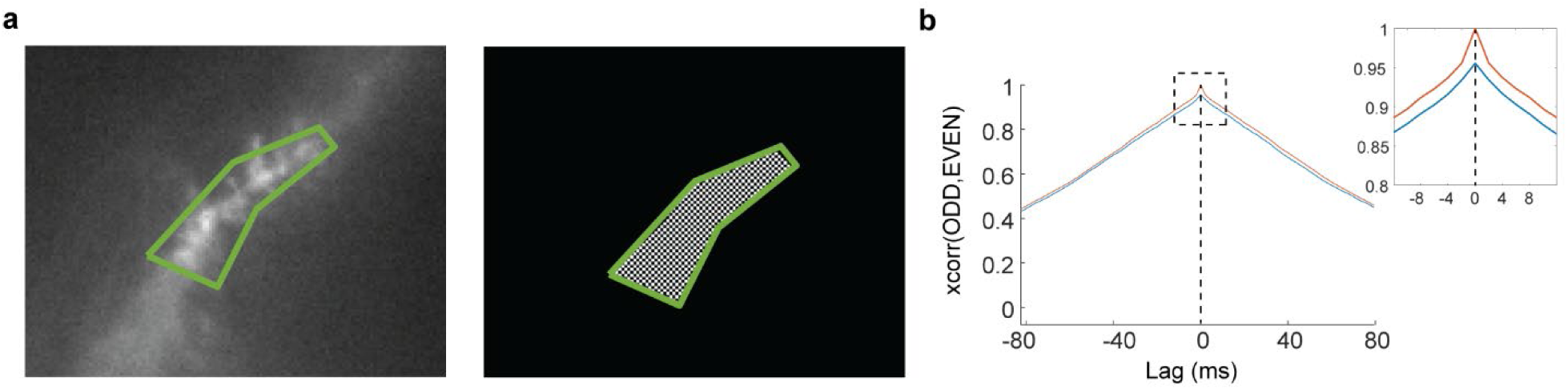
Contribution of shot noise to imperfect dendrite-to-dendrite cross-correlation. **a**, We isolate a region of interest (ROI) around each dendritic branch and split it into two signals derived from interleaved checkerboard patterns of pixels. These two signals are expected to be nearly identical, except for independent realizations of the photon shot noise. **b**, We compare the autocorrelation of the signal in the green ROI (orange plot) to the cross-correlation of the signals derived from the even and odd pixels (blue plot). The cross-correlation at zero-lag represents the maximum possible cross-correlation between dendritic branches, taking into account the contribution of shot noise.

**Figure S10.**
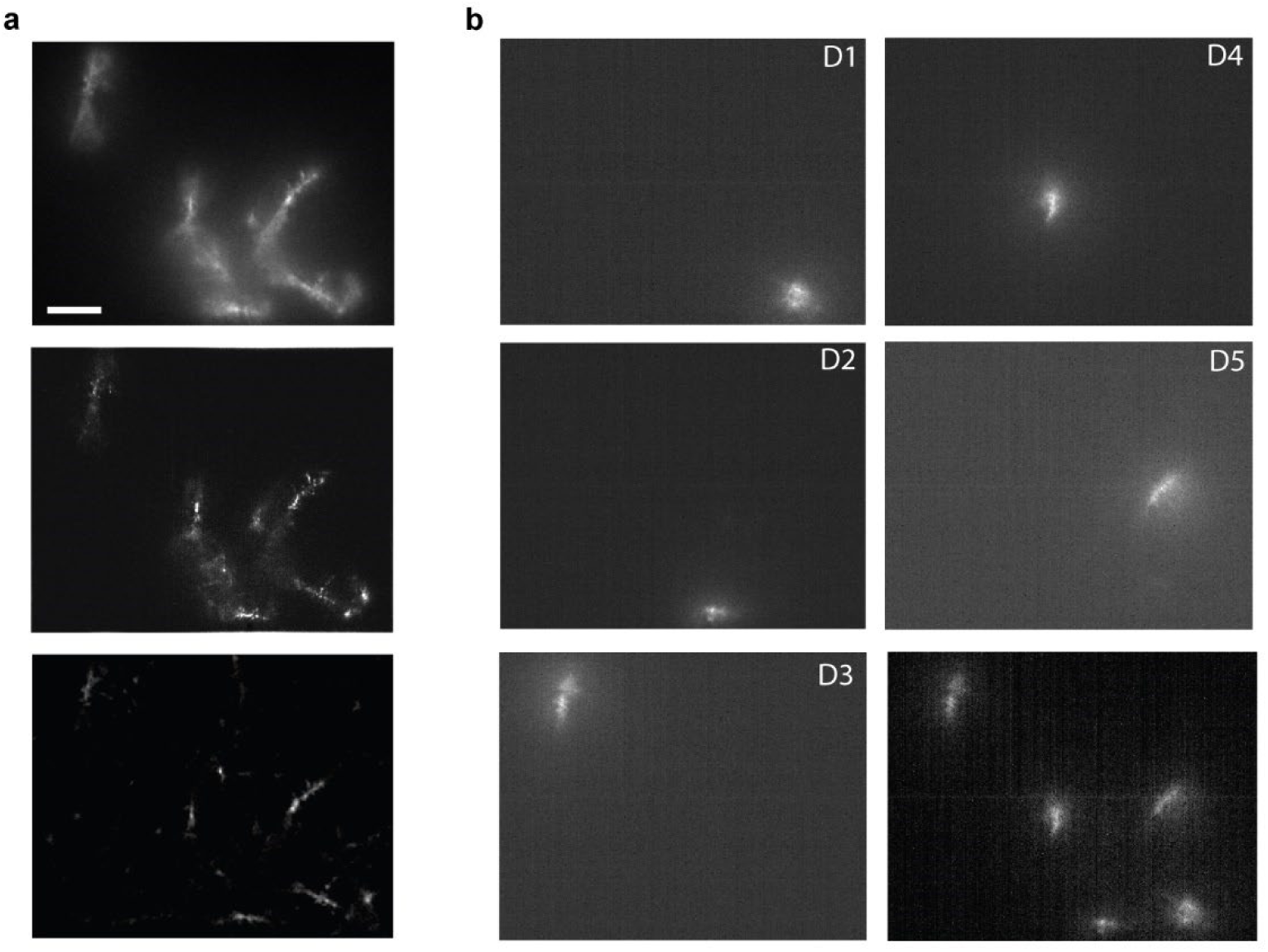
Localization of voltage imaging signals and optogenetic stimuli. **a**, Dendrites of neuron of **Fig. 3** imaged in a single focal plane. Top: Time-average fluorescence of Voltron2; Middle: 1-lag autocorrelation of a voltage imaging movie, showing localization of the fluctuating components of the signal; Bottom: HiLo image of the same field of view. Scale bar 20 μm. Scaling is the same for all figures. **b**, Each of five dendritic branches in (**a**) was sequentially targeted with optogenetic stimulation. The pattern of optogenetic stimulation was mapped via the fluorescence of eYFP in CheRiff-eYFP. Images D1 – D5 show eYFP fluorescence as the optogenetic stimulus was targeted to different branches. The absence of fluorescence from non-targeted branches confirms that the absence of optical crosstalk. Stimuli were kept localized to minimize spurious activation of non-targetted branches. Bottom right: Composite image in the eYFP channel showing the five stimulus patterns.

**Figure S11.**
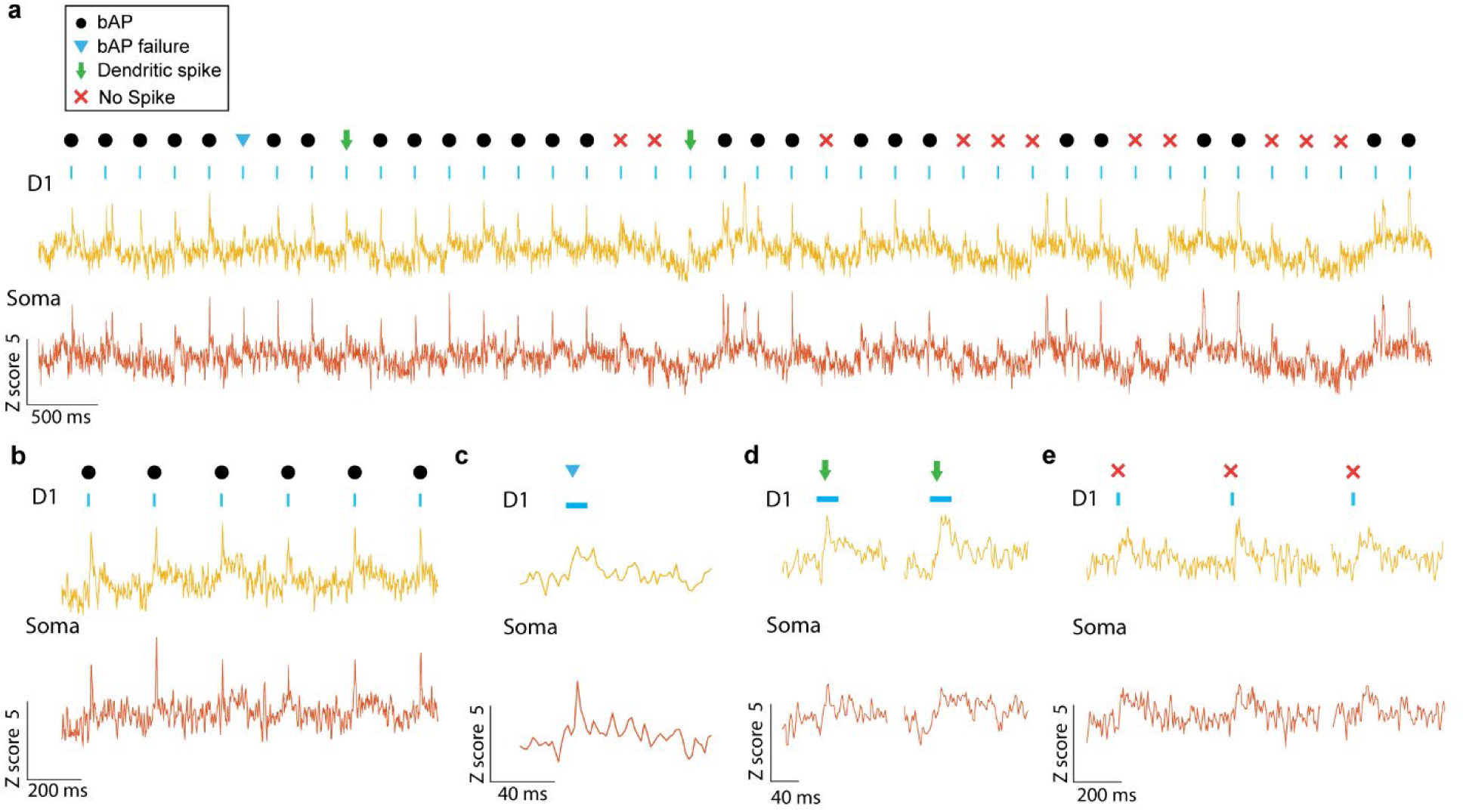
Response at dendrite and soma to strong dendrite-targeted stimuli. **a**, Example recording in which there is a good correspondence between spike-like events at the soma and dendrite under dendrite-targeted stimulation. **b**, In most cases, dendrite stimulation evoked spikes at the soma and dendrite. Other rare events comprised: **c**, Putative failed bAP. **d**, Putative dendritic spikes, comprising depolarization transients detected at the dendrites and not the soma. **e**, Examples where the stimulus triggered only subthreshold events at dendrite and soma. Stimuli that failed to evoke spikes at the soma or dendrite tended to coincide with subthreshold hyperpolarization.

**Figure S12.**
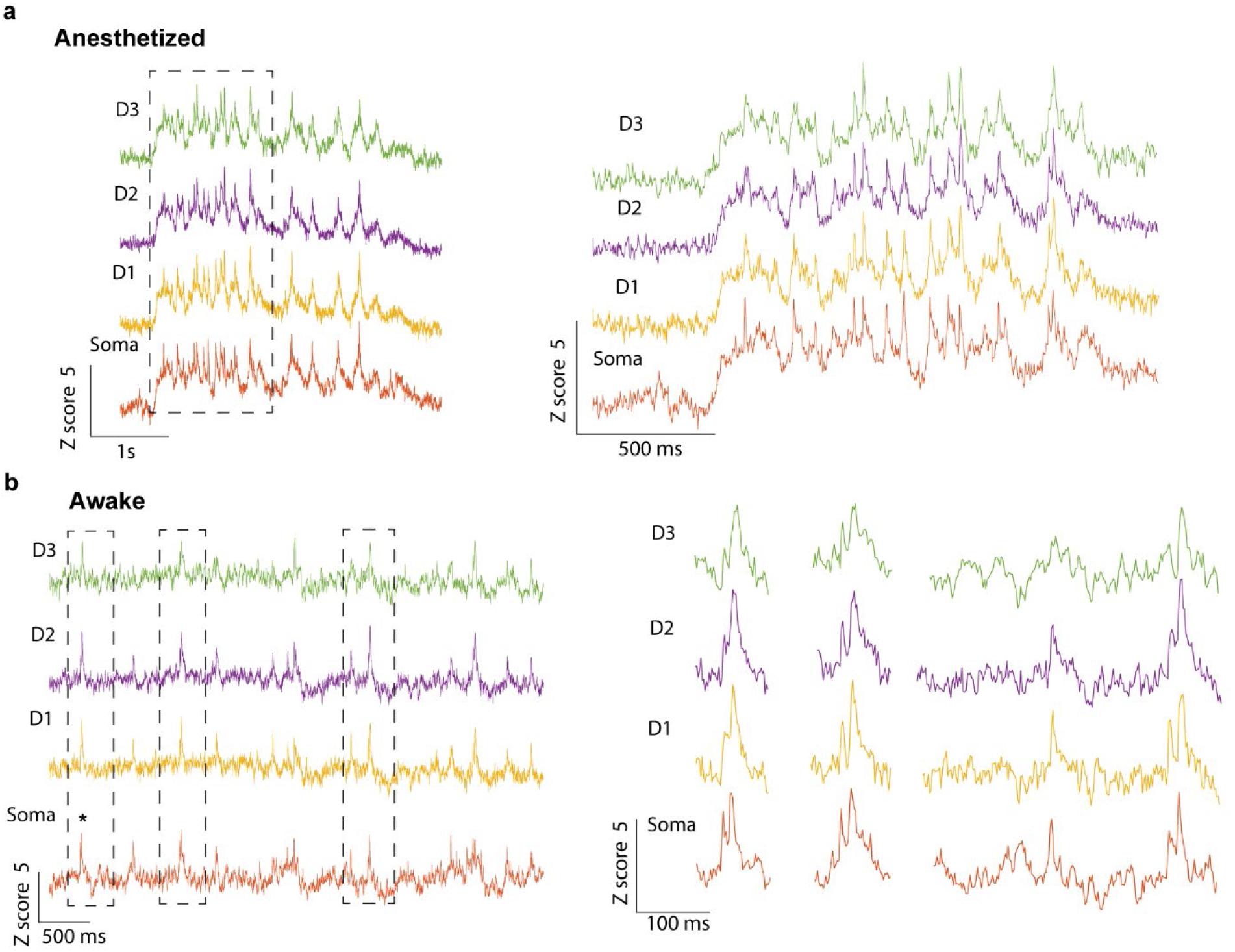
Spontaneous dynamics and bAP failures. Example traces of spontaneous activity for anesthetized (**a**) and awake (**b**). We observe bAP failures in both anesthetized and awake recordings.

**Figure S13.**
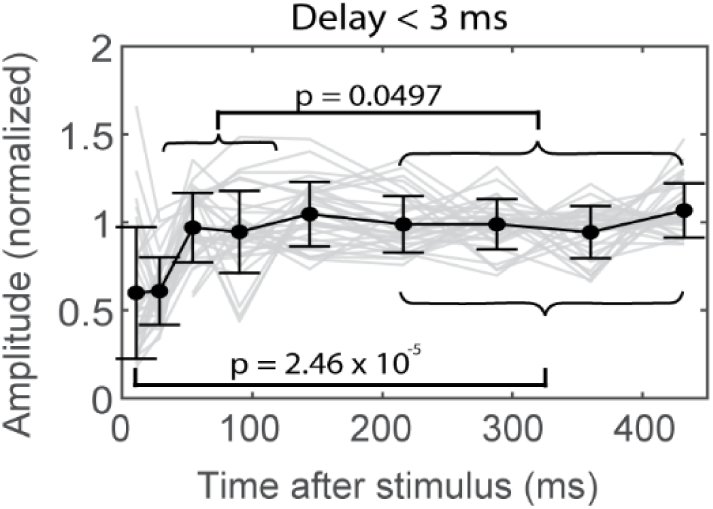
Dependence of bAP propagation on electrotonic distance. **a**, Dendritic branches closer to the soma (as determined by bAP delay of < 3 ms) showed less modulation of bAP amplitude compared to branches further from the soma (**Fig. 4e-h**).

**Figure S14.**
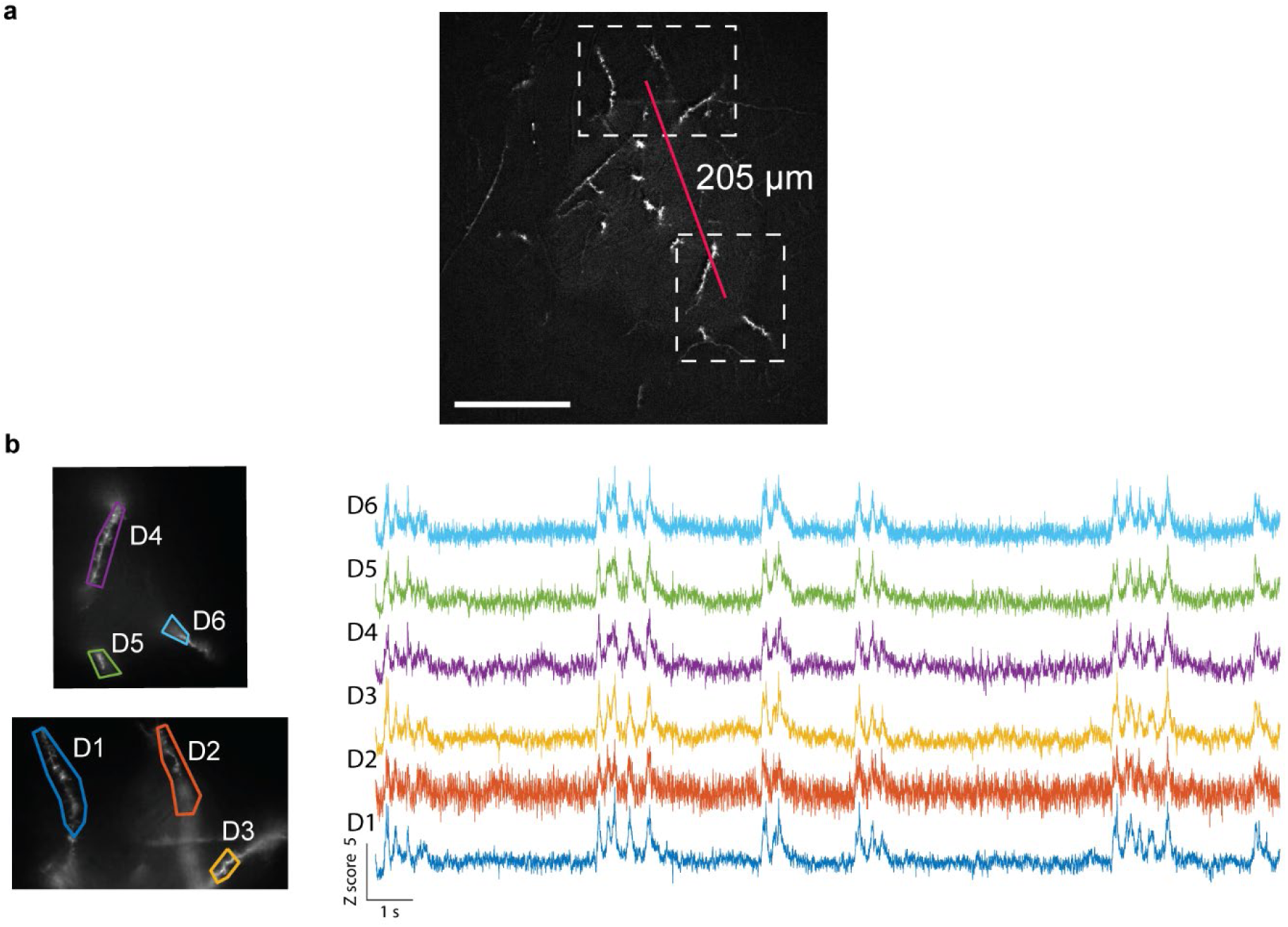
Recording dendritic voltages in two laterally offset fields of view. Independent control of the field of view of each camera allows simultaneous recording of distant branches from the same neuron. **a**, Wide-area HiLo image of an apical dendrite arbor. The two cameras were set to record from regions separated center-to-center by 205 μm. Scale bar 100 μm. **b**, Individual camera recording fields and single-dendrite regions of interest. **c**, Example traces from the two fields of view. We observed high correlation of up-down states across all recorded branches.

## References

1. Stuart, G., Spruston, N. & Häusser, M., Dendrites (Oxford University Press, 2016).

2. Spruston, N., Pyramidal neurons: dendritic structure and synaptic integration. Nature Reviews Neuroscience 9, 206–221 (2008).

3. London, M. & Häusser, M., Dendritic computation. Annual Review of Neuroscience 28, 503–532 (2005).

4. Häusser, M., Spruston, N. & Stuart, G. J., Diversity and Dynamics of Dendritic Signaling. Science 290, 739–744 (2000).

5. Magee, J. C. & Johnston, D., Plasticity of dendritic function. Current Opinion in Neurobiology 15, 334–342 (2005).

6. Stuart, G. J. & Spruston, N., Dendritic integration: 60 years of progress. Nature Neuroscience 18, 1713–1721 (2015).

7. Fischer, L. et al., Dendritic Mechanisms for In vivo Neural Computations and Behavior. The Journal of Neuroscience 42, 8460–8467 (2022).

8. Beaulieu-Laroche, L., Toloza, E. H. S., Brown, N. J. & Harnett, M. T., Widespread and Highly Correlated Somato-dendritic Activity in Cortical Layer 5 Neurons. Neuron 103, 235–241.e4 (2019).

9. Palmer, L. M. et al., NMDA spikes enhance action potential generation during sensory input. Nature Neuroscience 17, 383–390 (2014).

10. Jia, H., Rochefort, N. L., Chen, X. & Konnerth, A., Dendritic organization of sensory input to cortical neurons in vivo. Nature 464, 1307–1312 (2010).

11. O’Hare, J. K. et al., Compartment-specific tuning of dendritic feature selectivity by intracellular Ca^2+^ release. Science 375 (2022).

12. Gambino, F. et al., Sensory-evoked LTP driven by dendritic plateau potentials in vivo. Nature 515, 116–119 (2014).

13. Smith, S. L., Smith, I. T., Branco, T. & Häusser, M., Dendritic spikes enhance stimulus selectivity in cortical neurons in vivo. Nature 503, 115–120 (2013).

14. Waters, J. & Helmchen, F., Boosting of Action Potential Backpropagation by Neocortical Network Activity In Vivo. The Journal of Neuroscience 24, 11127–11136 (2004).

15. Margrie, T. W. et al., Targeted Whole-Cell Recordings in the Mammalian Brain In Vivo. Neuron 39, 911–918 (2003).

16. Kamondi, A., Acsády, L. & Buzsáki, G., Dendritic Spikes Are Enhanced by Cooperative Network Activity in the Intact Hippocampus. The Journal of Neuroscience 18, 3919–3928 (1998).

17. Abdelfattah, A. S. et al., Sensitivity optimization of a rhodopsin-based fluorescent voltage indicator. Neuron (2023).

18. Hochbaum, D. R. et al., All-optical electrophysiology in mammalian neurons using engineered microbial rhodopsins. Nature Methods 11, 825–833 (2014).

19. Shepard, B. D., Natarajan, N., Protzko, R. J., Acres, O. W. & Pluznick, J. L., A Cleavable N-Terminal Signal Peptide Promotes Widespread Olfactory Receptor Surface Expression in HEK293T Cells. PLoS ONE 8, e68758 (2013).

20. Lim, D., Chu, K. K. & Mertz, J., Wide-field fluorescence sectioning with hybrid speckle and uniform-illumination microscopy. Optics Letters 33, 1819 (2008).

21. Goetz, L., Roth, A. & Häusser, M., Active dendrites enable strong but sparse inputs to determine orientation selectivity. Proceedings of the National Academy of Sciences 118 (2021).

22. Boahen, K., Dendrocentric learning for synthetic intelligence. Nature 612, 43–50 (2022).

23. Petersen, C. C. H., Hahn, T. T. G., Mehta, M., Grinvald, A. & Sakmann, B., Interaction of sensory responses with spontaneous depolarization in layer 2/3 barrel cortex. Proceedings of the National Academy of Sciences 100, 13638–13643 (2003).

24. Waters, J. & Helmchen, F., Background Synaptic Activity Is Sparse in Neocortex. The Journal of Neuroscience 26, 8267–8277 (2006).

25. P Park, e. a., Dendritic voltage imaging reveals biophysical basis of associative plasticity rules. BioArxiv (2023).

26. Hoffman, D. A., Magee, J. C., Colbert, C. M. & Johnston, D., K+ channel regulation of signal propagation in dendrites of hippocampal pyramidal neurons. Nature 387, 869–875 (1997).

27. Jung, H.-Y., Mickus, T. & Spruston, N., Prolonged Sodium Channel Inactivation Contributes to Dendritic Action Potential Attenuation in Hippocampal Pyramidal Neurons. The Journal of Neuroscience 17, 6639–6646 (1997).

28. Payeur, A., Guerguiev, J., Zenke, F., Richards, B. A. & Naud, R., Burst-dependent synaptic plasticity can coordinate learning in hierarchical circuits. Nature Neuroscience 24, 1010–1019 (2021).

29. Landau, A. T. et al., Dendritic branch structure compartmentalizes voltage-dependent calcium influx in cortical layer 2/3 pyramidal cells. eLife 11 (2022).

30. Adam, Y. et al., Voltage imaging and optogenetics reveal behaviour-dependent changes in hippocampal dynamics. Nature 569, 413–417 (2019).

31. Goldey, G. J. et al., Removable cranial windows for long-term imaging in awake mice. Nature Protocols 9, 2515–2538 (2014).

32. Waters, J., Sources of widefield fluorescence from the brain. eLife 9 (2020).

33. Lin, D. et al., Time-tagged ticker tapes for intracellular recordings. Nature Biotechnology 41, 631–639 (2023).

